# Auranofin Inhibits Hepatitis E Virus Replication Via Reactive Oxygen Species

**DOI:** 10.1101/2025.05.02.651833

**Authors:** Kateland Tiller, S. Tyler Williams, Bo Wang, Debin Tian, Xiang-Jin Meng, James Weger-Lucarelli

## Abstract

Hepatitis E virus (HEV) causes roughly 20 million yearly global infections, and is associated with chronic hepatitis, neurological sequelae and pregnancy-related adverse outcomes that require antiviral therapeutic intervention. While there are currently no approved HEV-specific therapeutics, ribavirin and pegylated interferon, prescribed off-label, are the current standard of care. However, ribavirin resistance and toxicity highlight the unmet clinical need to identify safer, HEV-specific antivirals. Auranofin, an FDA-approved anti-rheumatic drug, displays antiviral activity against several viruses. Therefore, we investigated auranofin’s potential as an antiviral and its mechanism of action against HEV. We demonstrated that auranofin displays dose-dependent antiviral activity against two genotypes of HEV that cause a significant proportion of human disease, as well as against a ribavirin treatment failure-associated mutant. Because auranofin is known to increase reactive oxygen species (ROS), we investigated the antiviral mechanism of action via treatment with ROS inhibitors. ROS inhibitors reversed auranofin-mediated ROS promotion and antiviral activity, suggesting the observed antiviral effects are mediated by ROS. Furthermore, treatment with a different ROS promotor, D-amino acid oxidase (DAAO), also displays antiviral activity against HEV, which was also reversed by treatment with a ROS inhibitor, suggesting that ROS accumulation alone is antiviral. We also demonstrated that combined treatment with auranofin and ribavirin exhibits synergistic antiviral activity *in vitro*, which supports repurposing auranofin as an antiviral against HEV, potentially in combination with ribavirin. Overall, this study has important implications in repurposing auranofin as an antiviral against HEV and in delineating the mechanism of action against HEV via ROS.

**Importance:** Hepatitis E virus (HEV) lacks approved antiviral therapies, and off-label treatments are limited by toxicity and emerging resistance. This study identifies the FDA-approved drug auranofin as an effective in vitro inhibitor of HEV, including two globally relevant human-associated genotypes and a ribavirin treatment failure-associated mutant. Auranofin’s activity highlights the therapeutic potential of host-targeting antivirals, particularly those that promote the generation of reactive oxygen species, in treating HEV infection. These findings support further in vivo investigations of auranofin as a treatment for HEV and suggest that modulating host redox pathways by promoting reactive oxygen species may represent a promising strategy for broad-spectrum antiviral development.

## Introduction

Hepatitis E virus (HEV) is a globally distributed virus that has been estimated by the World Health Organization (WHO) to cause approximately 20 million global infections every year, with 3.3 million symptomatic infections and approximately 44,000 deaths, making HEV a major cause of acute and chronic viral hepatitis worldwide (1–3). Most HEV infections in developing countries are acquired through virus-contaminated water during large-scale outbreaks. In contrast, in industrialized countries with good sanitary infrastructure, sporadic cases often result from zoonotic transmission through consumption of undercooked or raw meat products (4, 5). Based on these multiple routes of transmission, it is suggested that the actual global disease burden caused by HEV is greatly underestimated (6, 7).

HEV is a single-stranded positive-sense RNA virus belonging to the family *Hepeviridae*, which consists of two subfamilies: *Orthohepevirinae* and *Parahepevirinae* (8). The major HEV genotypes known to infect humans belong to the species *balayani* in the genus *Paslahepevirus*. Genotypes 1 and 2 (HEV-1 and HEV-2) exclusively infect humans and are responsible for large-scale outbreaks in developing countries. In contrast, HEV-3 and HEV-4 infect both humans and other animals, causing sporadic cases of zoonotic transmission (9). HEV is a non-enveloped, spherical virus with particles of approximately 30-35 nm in stool samples, although virions circulating in the blood of infected individuals and those produced through cell culture exist as quasi-enveloped particles (10–13). The HEV genome is approximately 7.2 kb in length, containing three partially overlapping open reading frames (ORFs): ORF1 encodes non-structural proteins responsible for replication, ORF2 encodes the structural capsid protein, and ORF3 encodes a small protein involved in virus replication and assembly (14–17).

Beyond its virological characteristics, HEV poses a significant global health burden due to its varied clinical manifestations and potential for severe disease in vulnerable populations. While most patients experience a self-limiting, acute infection, HEV infection can also result in chronic and/or deadly outcomes in at-risk groups, including immunocompromised people, pregnant women, or those who have pre-existing liver disease (18–20). Immunocompromised individuals, such as those who are HIV-positive, solid organ transplant recipients, and patients undergoing chemotherapy treatments, are more likely to develop chronic HEV infections, which can become deadly if liver fibrosis or cirrhosis develops (21–23). Severe disease outcomes associated with HEV infection disproportionately affect pregnant women and their developing fetuses. HEV infection in the second or third trimesters of pregnancy greatly increases the risk of developing fulminant hepatic failure (FHF) and death (19, 24). Vertical transmission has also been reported, resulting in adverse fetal outcomes of preterm delivery, fetal distress, and/or low birth weight (25), although others failed to transmit HEV vertically under experimental conditions (26, 27). A significant proportion of HEV-infected individuals develop various neurological sequelae, including Guillain-Barré syndrome and neuralgic amyotrophy (28–30). Recently, HEV has been recognized as the third leading cause of foodborne viral illness (31), and pork is a leading source of foodborne HEV infections (32–34). Chronic hepatitis E, HEV-associated neurological complications, foodborne hepatitis E, and the severity of disease outcomes in certain at-risk populations highlight the urgent need for effective antivirals against HEV.

Currently, no therapeutics are approved for treating hepatitis E, and the only hepatitis E vaccine is approved for use in China (35) and Pakistan (36). Ribavirin and pegylated interferon are used as off-label treatments and represent the current standard of care for treating HEV infection. However, ribavirin resistance has been reported (37, 38), and pegylated interferon is associated with significant side effects (39–41), thus highlighting the need for more effective and HEV-specific antivirals to treat hepatitis E.

Recently, there has been an increasing interest in repurposing FDA-approved drugs as antivirals. This process is faster, safer, and more cost-effective than identifying and developing novel antivirals (42, 43). One such FDA-approved drug is auranofin, a gold-based compound used in patients with rheumatoid arthritis. Auranofin has shown promise in treating a wide range of ailments, including viral, bacterial, and fungal infections as well as cancer (44). Notably, it has demonstrated antiviral activity against chikungunya virus (44), human immunodeficiency virus (45, 46), and SARS-CoV-2 (47, 48). Because auranofin displays broad-spectrum antiviral activity, we sought to test its potential antiviral activity against HEV. We tested the antiviral activity of auranofin against different relevant strains of human HEV in a human hepatocyte cell line. We found that auranofin displayed dose-dependent antiviral activity against HEV at non-toxic concentrations. Further studies into the antiviral mechanism of action revealed that reactive oxygen species (ROS) mediate auranofin’s antiviral activity, and that promoting ROS alone displayed robust antiviral activity against HEV. We also showed that combined treatment with ribavirin and auranofin provides a synergistic antiviral effect. Collectively, our results suggest that auranofin is a promising antiviral against HEV infection and may be used off-label in conjunction with ribavirin to treat HEV infections, particularly chronic hepatitis E and HEV-associated neurological sequelae.

## Methods and Materials

### Cell culture

Huh7-S10-3 cells, a sub-clone of human hepatocyte cellular carcinoma cells (Huh7) (49), kindly provided by Suzanne U. Emerson (NIAID, NIH, Bethesda, MD), were cultured in Dulbecco’s modified Eagle’s medium (DMEM) (Corning) with high glucose, L-glutamine, and sodium pyruvate supplemented with 10% fetal bovine serum (FBS), 1% non-essential amino acids, 0.1% gentamicin sulfate, and 25 mM HEPES (herein called DMEM-10). The cells were incubated at 37 °C with 5% CO_2_.

### Compounds

Auranofin, kindly provided by Veronica Ghini (Resonance Magnetic Center, Italy), was prepared in dimethyl sulfoxide (DMSO) (Sigma Aldrich) to a stock concentration of 10 mM. Additional auranofin stocks (Selleckchem) were also prepared to 10 mM in DMSO. Ribavirin (Thermofisher Scientific) was prepared in molecular grade water to a concentration of 40.95 mM. N-acetylcysteine (NAC) (Thermo Fisher Scientific) was prepared fresh in molecular-grade water to varying concentrations at a pH of 8 with the addition of sodium hydroxide (NaOH). Dithiothreitol (DTT) (VWR) was purchased at a concentration of 1 M. D-amino acid oxidase (DAAO) (Millipore-Sigma) was prepared in water to a concentration of 22 mg/mL.

### HEV indicator replicons and infectious clones

The HEV-1 Sar55 *Gaussia* luciferase (Gluc) indicator replicon was constructed using the HEV-1 Sar55 strain backbone (GenBank accession no. AF444002) (50), in which a portion of ORF2 was replaced with the *Gaussia* luciferase gene (51). Similarly, the HEV-3 P6 Gluc indicator replicon was developed from the HEV-3 strain Kernow-C1/P6 (designated as P6) backbone (GenBank accession no. JQ679013) that was serially passaged six times in cell culture (52, 53). Additionally, a G1634R mutation was introduced into the HEV-3 P6 Gluc construct, based on prior findings that show this mutation significantly enhances HEV-3 replication *in vitro* and promotes ribavirin resistance *in vivo* (54). This mutant was constructed using a site-directed mutagenesis system, as previously described (55). A HEV-1 infectious luminescence reporter virus Sar55(Hib), which has a HiBiT tag in the C-terminal of ORF2 of the wild-type Sar55 strain, was generated as previously described (56). The HEV-3 P6 Gluc indicator replicon was kindly provided by Dr. Sue Emerson of the National Institute of Allergy and Infectious Diseases at NIH. The HEV-1 Sar55 Gluc indicator replicon was generously provided by Dr. Alexander Ploss of Princeton University, Princeton, NJ.

### Replicon and infectious clone amplification, *in vitro* transcription, and transfection

Replicon or infectious clone plasmids were amplified via rolling circle amplification (RCA) as previously described (57). The RCA product was then linearized with the restriction enzyme MluI (for P6-based) or BglII (for Sar55-based) (New England Biolabs). The linearized product was purified using SPARQ PureMag Beads (Quantabio), and *in vitro* transcription was performed with the mMESSAGE mMACHINE T7 Transcription Kit (Thermofisher Scientific) to produce capped, infectious RNA. Viral RNA was transfected via the jetMESSENGER mRNA transfection reagent (Polyplus) into Huh7-S10-3 cells and incubated with 5% CO_2_ at 37 °C.

### Cytotoxicity assay

Huh7-S10-3 cells were plated at 5,000 cells/well in a 96-well plate and incubated at 37 °C with 5% CO_2_. Compound dilutions were prepared in DMEM-10. The vehicle (DMSO or H_2_O) was also prepared to corresponding concentrations to serve as a control. Growth media was removed from the cells and replaced with the same volume of each compound dilution. Cells were incubated for 72 hours before CellTiter 96 AQueous One Solution Reagent was added to each well as recommended by the manufacturer’s protocol (Promega CellTiter 96 AQueous One Solution Cell Proliferation Assay). Plates were incubated at 37 °C with 5% CO_2_ for 2-4 hours. After incubation, the absorbance was measured at 490 nm using an Infinite M plate reader. Viability was calculated by subtracting media only absorbance values and normalizing to the appropriate vehicle control. Cytotoxic concentration 50% (CC_50_) values were generated via the ED50 Plus v1.0 software (58).

### HEV antiviral assays

Huh7-S10-3 cells were plated at 5,000 cells/well in a 96-well plate. Cells were transfected with HEV replicon- or infectious clone-derived RNA as described above. After a 4-hour incubation, compound dilutions were prepared in DMEM-10. The vehicle (DMSO or H_2_O) was also prepared to a corresponding concentration to serve as a control. Growth media was removed from the cells and replaced with the same volume of each compound dilution. For the HEV-3 P6 Gluc replicon testing, cells were incubated with the compounds for 72 hours. For HEV-1 Sar55 Gluc replicon and infectious reporter virus Sar55(Hib) testing, cells were incubated with the compounds for 7 days. *Gaussia* luciferase expression was quantified from the supernatant via the Pierce™ *Gaussia* luciferase glow assay kit (Thermofisher Scientific). HiBiT activity in the supernatant of HEV-1 Sar55(Hib) transfected cells was quantified using the Nano-Glo® HiBiT Extracellular Detection System (Promega). Luminescence was measured on an Infinite M plex multimode microplate reader for both Gluc and HiBiT activities. Antiviral activity was calculated by subtracting the negative transfection only luminescence values and normalizing to the appropriate vehicle control values. The half maximal effective concentration (EC_50_) values were generated via ED50 Plus v1.0 software (58) for select experiments.

### ROS quantification

Huh7-S10-3 cells were plated in 24-well plates to 50,000 cells/well. Compounds were prepared in DMEM-10 to the desired concentrations. The 2’,7’-dichlorodihydrofluorescein diacetate (H_2_DCFDA) (Fisher Scientific) probe was also added to each compound dilution at a final concentration of 10 μM. Compound dilutions were then applied to the cells, and the plates were incubated for 30 minutes at 37 °C with 5% CO_2_. The cells were then trypsinized, and DMEM-10 was added to create a single-cell suspension. Cells from two wells were pooled for each sample and pelleted at 300×g for 5 min at 4°C. Then, the cells were washed with PBS, re-pelleted, and finally resuspended in 100 μL of PBS. Samples were analyzed by flow cytometry using the FACSAria Fusion Flow cytometer (BD Biosciences), and the median fluorescent peak was recorded. The data was then analyzed by normalization to the appropriate vehicle control.

### Synergy analysis

Huh7-S10-3 cells were plated at 5,000 cells/well in a 96-well plate. Cells were transfected with HEV-3 P6 Gluc RNA, and compounds were prepared in DMEM-10 to the desired concentrations. Auranofin was tested at concentrations of 0, 0.5, 1, and 1.5 μM, and ribavirin was tested at concentrations of 0, 5, 10, 15, 20, and 25 μM. Each concentration was tested alone and in combination with auranofin and ribavirin, with all possible concentration combinations being tested. Compounds were applied, and 72 hours later, Gluc expression was quantified from the supernatant. Antiviral activity was calculated by subtracting the negative transfection only luminescence values and normalizing to the appropriate vehicle control values. Data was uploaded to the SynergyFinder web application (version 3.0) (59). The synergy between auranofin and ribavirin was determined using the highest single agent (HSA) model. This model determines if the combined effect of two drugs exceeds the sum of their individual effects; if so, then a synergistic effect exists. Combinations determined to have a synergy score greater than 10 were graphed independently for statistical analysis.

### Statistical analysis

Statistical analyses were performed using Graphpad Prism 10. For initial cytotoxicity and antiviral testing, tested compound concentrations were converted to a log scale and a non-linear regression ([log inhibitor] vs. normalized response) was performed. To determine EC_50_ and CC_50_ values, data was generated via ED50 Plus v1.0 software (58). For all other antiviral and ROS testing, data was analyzed via one-way analysis of variance (ANOVA) using Dunnett’s test for multiple comparisons. Combined testing of auranofin and ribavirin was analyzed via the SynergyFinder web application (version 3.0) (59). Select compound combinations were analyzed via one-way ANOVA as previously mentioned.

## Results

### Auranofin displays dose-dependent antiviral activity against HEV-1, Sar55 Gluc replicon, and HEV-1 infectious reporter virus Sar55(Hib)

To establish a range of non-toxic concentrations to test auranofin’s antiviral activity, cell viability was assessed 72 hours post-compound application on Huh7-S10-3 cells via an MTS assay. At concentrations below 2 μM, auranofin displayed low toxicity in Huh7-S10-3 cells. However, cell viability decreased at concentrations above 2 μM, resulting in a CC_50_ of 2.53 μM (Fig. 1A). Based on these results, we performed subsequent experiments using auranofin at concentrations of 2 μM or below.

**Figure 1.**
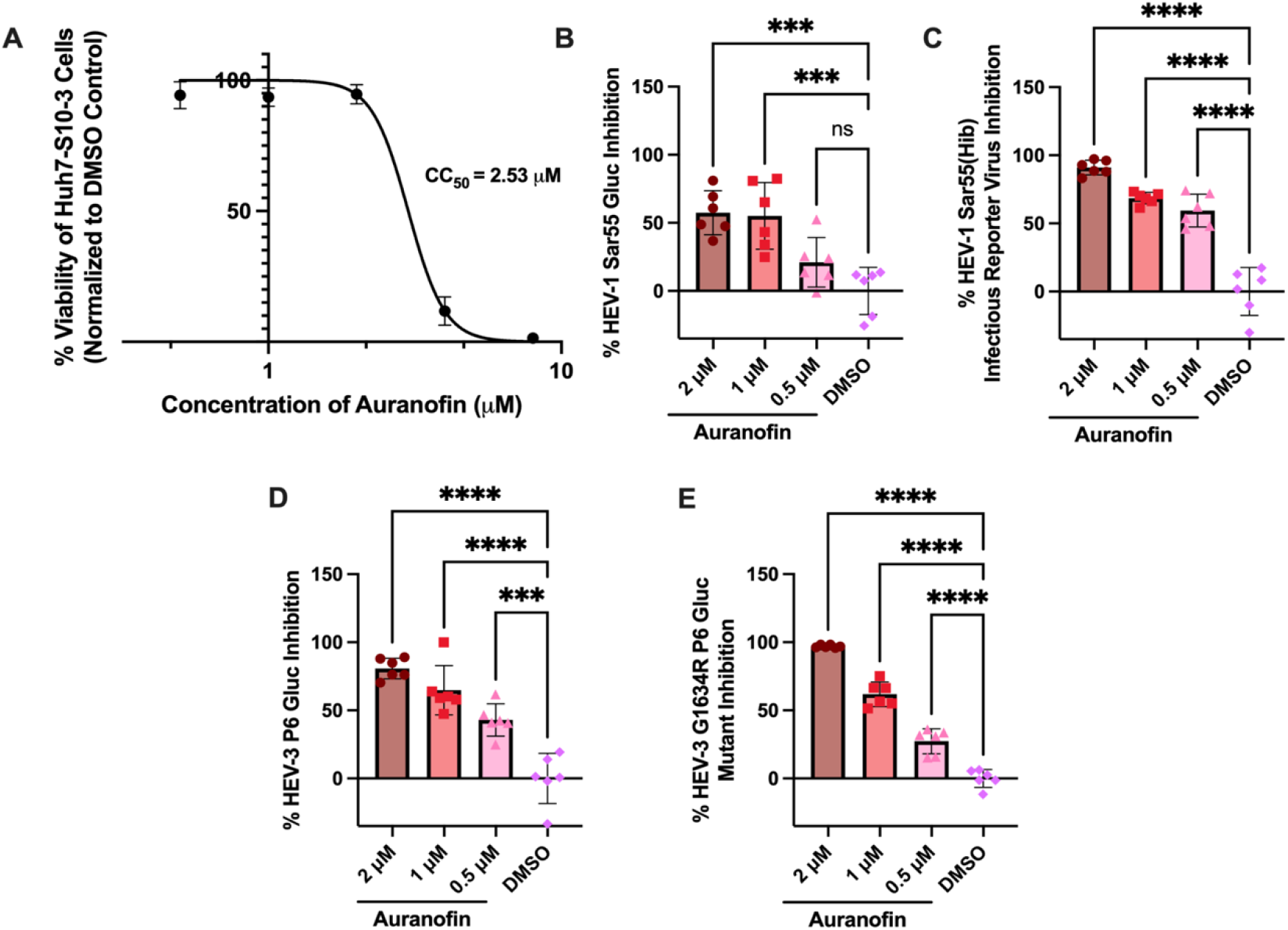
Auranofin displays antiviral activity in a dose-dependent manner at non-toxic concentrations against two different genotypes of HEV and a ribavirin treatment failure-associated HEV mutant. (*A*) The effect of auranofin on cell viability in Huh7-S10-3 (human hepatocyte) cells, measured via MTS assay. Data was collected 72 hours after addition of the compound, background subtracted and normalized to the DMSO vehicle control. X-axis values are log transformed. The lines represent non-linear regression curves, and data points represent means ± SD; n=6 performed in two independent experiments. (*B & C*) Antiviral activity of auranofin against the HEV-1 Sar55 Gluc replicon (*B*) and HEV-1 infectious reporter virus Sar55 (Hib) (*C*). (*D & E*) Antiviral activity of auranofin against the HEV-3 P6 Gluc replicon (*D*) and the HEV-3 G1634R P6 gluc ribavirin resistance mutant (*E*). For antiviral assays, data was collected 72 hours post-transfection for Gluc replicons and 7 days post-transfection for Sar55(Hib) infectious reporter virus. Data from B-E was background subtracted, normalized to a DMSO concentration corresponding to that present in the 2 μM auranofin group, and was analyzed via one-way ANOVA. Error bars represent ± SD; n=6 performed in two separate experiments.

To thoroughly analyze auranofin’s potential antiviral activity against HEV, we first tested non-toxic doses against the HEV-1 Sar55 *Gaussia* luciferase (Gluc) replicon and the HEV-1 Sar55(Hib) HiBiT-tagged infectious reporter virus that enables luciferase-based detection of extracellular viral particles (56). The Gluc expression (72 h post-inoculation) and Hibit expression (7 days post-inoculation) were measured to indicate the replication of HEV-1. We showed that auranofin has dose-dependent antiviral activity against the HEV-1 Sar55 Gluc replicon (Fig. 1B) and the HEV-1 Sar55(Hib) HiBiT-tagged infectious virus (Fig. 1C), demonstrating auranofin’s antiviral activity against a replicon and infectious virus of HEV-1.

### Validation of auranofin’s antiviral activities using a different HEV genotype, HEV-3 P6 Gluc and a ribavirin treatment failure-associated HEV-3 P6 G1634R mutant

To determine if auranofin displays antiviral activity against other HEV genotypes, we tested the antiviral activity of auranofin by using a genotype 3 HEV (HEV-3 P6 Gluc replicon). We showed that auranofin displayed dose-dependent antiviral activity at non-toxic concentrations, resulting in an EC_50_ of 0.54 μM (Fig. 1D) (Supplementary Figure 1). Antiviral testing was also performed against a HEV-3 P6 Gluc mutant containing the G1634R mutation, which is associated with ribavirin treatment failure *in vivo*, to determine if auranofin is effective against a ribavirin treatment failure-associated HEV strain (37, 54). Similar to the WT HEV-3 P6 Gluc, the HEV-3 G1634R P6 Gluc mutant displayed similar susceptibility to auranofin treatment (Fig. 1E). Altogether, this *in vitro* data suggests that auranofin is an effective antiviral against two different genotypes of HEV and against a mutant associated with ribavirin treatment failure.

### ROS inhibitors reverse auranofin’s antiviral activity

Auranofin has been reported to inhibit glutathione peroxidase (GPx) and thioredoxin reductase (TrxR), resulting in increased intracellular ROS levels (60–63). Thus, we hypothesized that this ROS production may contribute to auranofin’s antiviral activity against HEV. To test this hypothesis, we treated cells with common ROS inhibitors, N-acetylcysteine (NAC) and dithiothreitol (DTT), in the presence of auranofin. NAC reduces ROS by promoting glutathione production, direct ROS scavenging, and modulating redox-signaling pathways (64, 65). DTT is thought to neutralize ROS and other free radicals, in addition to protecting against mitochondrial oxidative damage and regenerating glutathione from oxidized glutathione (GSSG) (66, 67), suggestive of differential anti-ROS mechanisms for NAC and DTT.

Co-treatment with 10 mM NAC and 2 μM auranofin completely reversed the antiviral activity of 2 μM auranofin against the HEV-3 P6 Gluc replicon (Fig. 2A). To further validate the impact on intracellular ROS levels, we used the fluorescent probe H_2_DCFDA to measure general intracellular ROS levels. When compounds and the H_2_DCFDA probe were applied to Huh7-S10-3 cells, we observed an increase in ROS in the presence of 2 μM auranofin (Fig. 2B). This increase was reversed with the combined treatment of 10 mM NAC, mirroring the reversal of antiviral activity. NAC treatment resulted in a robust reversal of baseline ROS below the DMSO control, to which the data were normalized, resulting in negative values. To test the contribution of ROS in an orthogonal manner, we also tested 500 μM DTT in the presence of 2 μM auranofin. Again, we observed a reversal of antiviral activity (Fig. 2C) and ROS promotion (Fig. 2D) in the presence of DTT. Together, these results suggest that ROS mediates auranofin’s antiviral effects against HEV.

**Figure 2.**
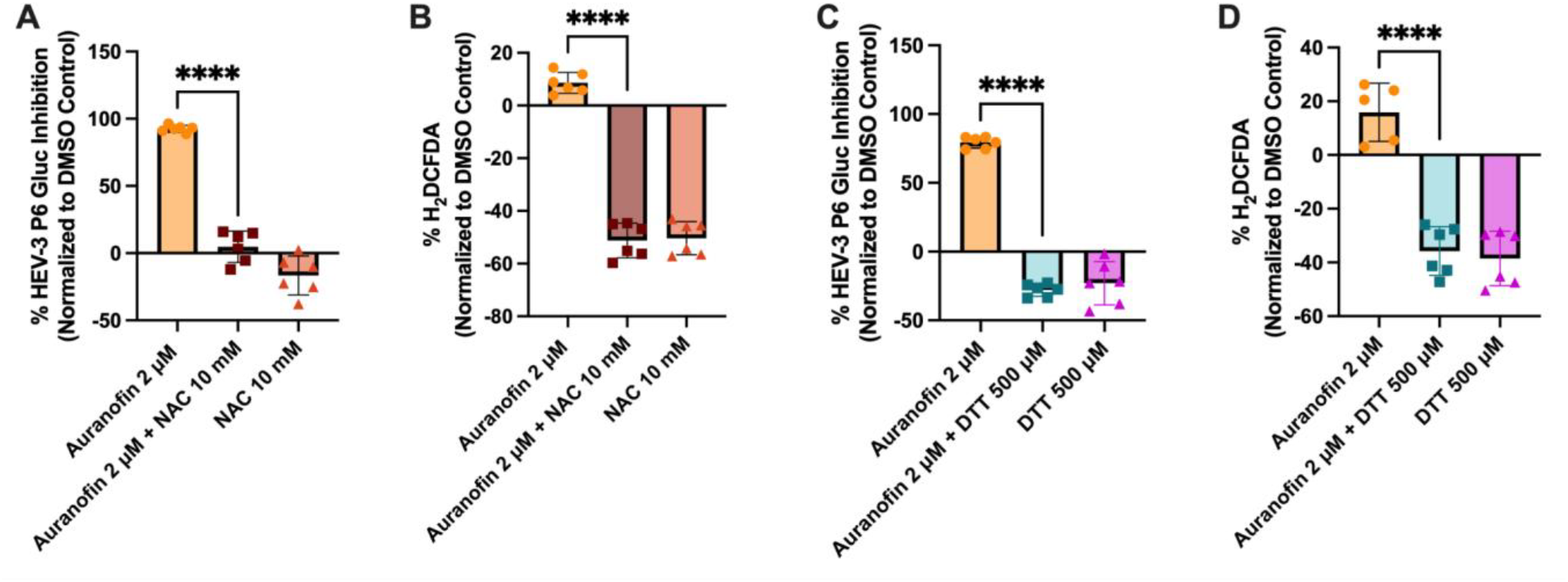
ROS inhibitors, NAC and DTT, reverse auranofin’s antiviral activity and ROS promotion. (*A & C*) The effect of 2 μM auranofin in the presence of 10 mM NAC (*A*) or 500 μM DTT (*C*) on HEV-3 P6 Gluc production as a proxy for HEV-3 P6 replication 72 hours post compound application. (*B & D*) The effect of 2 μM auranofin in the presence of 10 mM NAC (*B*) or 500 μM DTT (*D*) on general intracellular ROS via levels 10 μM treatment of H_2_DCFDA fluorescent probe and compounds of interest with a 30-minute incubation period. Single-cell suspensions were analyzed by flow cytometry, and the median fluorescent peak was recorded. Data is background subtracted and normalized to DMSO vehicle control. A one-way ANOVA analysis was performed. Data points represent means ± SD; n=6 performed in two independent experiments.

### The ROS promoter, D-amino acid oxidase (DAAO), displays antiviral activity against HEV

To further examine the antiviral effects of ROS, we tested the ROS promoter, D-amino acid oxidase (DAAO). DAAO has been shown to promote hydrogen peroxide (H_2_O_2_) production through the breakdown of D-amino acids, a ROS-promoting mechanism distinct from auranofin and independent of GPx and TrxR inhibition (68). DAAO displayed strong antiviral activity against HEV-3 P6 Gluc and no cytotoxicity at the doses tested, with a CC_50_ of >1000 μg/mL and an EC_50_ of 388.1 μg/mL (Fig. 3A). The antiviral activity (Fig. 3B) and ROS promotion (Fig. 3C) of 200 μg/mL DAAO were also reversed by co-treatment with 30 mM NAC. This data suggests that ROS promotion, independent of GPx and TrxR inhibition, inhibits HEV replication.

**Figure 3.**
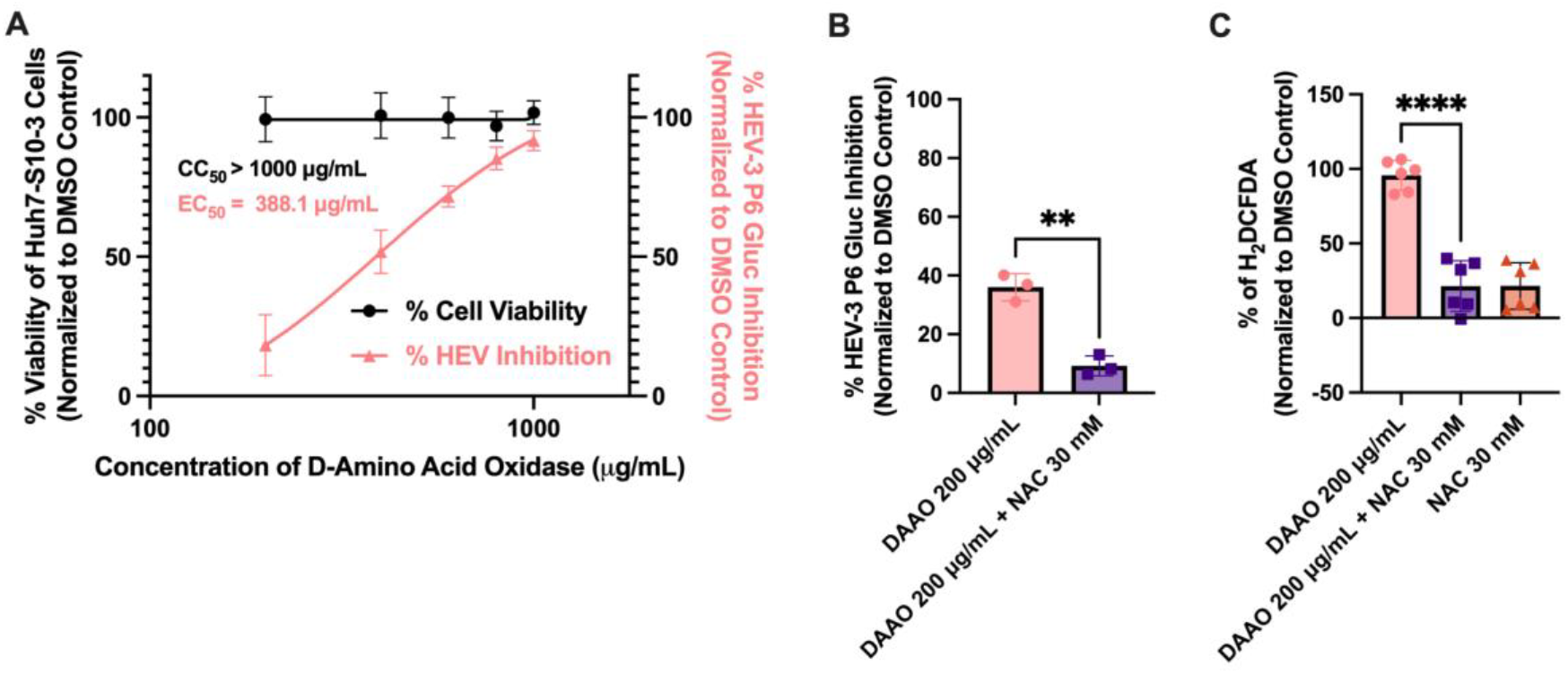
DAAO, a ROS promoter, displays antiviral activity, which is reversed by NAC, a ROS inhibitor. (*A*) The effect of DAAO on cell viability and antiviral activity of the HEV-3 P6 Gluc replicon in Huh7-S10-3 cells. Data collected 72 hours post compound application was background subtracted and normalized to the DMSO control. X-axis values are log transformed. The lines represent non-linear regression curves, and data points represent means ± SD; n=6 performed in two independent experiments. (*B*) The antiviral effect of 200 μg/mL DAAO in the presence of 30 mM NAC on the HEV-3 P6 Gluc replicon. (*C*) Generalized ROS measurement after 30-minute incubation with 10 μM treatment of H_2_DCFDA fluorescence probe. Samples were also incubated for 30 minutes with 200 μg/mL DAAO and 30mM NAC. Single-cell suspensions were analyzed via flow cytometry, and median fluorescent peak was recorded. Data from B and C are background-subtracted, normalized to DMSO, and analyzed via one-way ANOVA, with data points representing means ± SD. n = 6, performed in two independent experiments.

### Auranofin and ribavirin combined treatment exhibits synergistic antiviral effects *in vitro*

Since the antiviral activity of auranofin and ribavirin is likely through different mechanisms, we hypothesized that a combined treatment would exert synergistic antiviral activity (69). This is an important consideration because multiple ribavirin resistance mutants have been identified in patients, and a combined antiviral treatment could prevent the development of and further expansion of ribavirin-resistant HEV strains (70). We first assessed the cytotoxicity and antiviral activity of ribavirin to determine the optimal concentration range for combined treatment testing. Ribavirin displays stable cell viability in Huh7-S10-3 cells with a CC_50_ > 500 μM, and strong dose-dependent antiviral activity with an EC_50_ of 15.41 μM (Fig. 4A).

**Figure 4.**
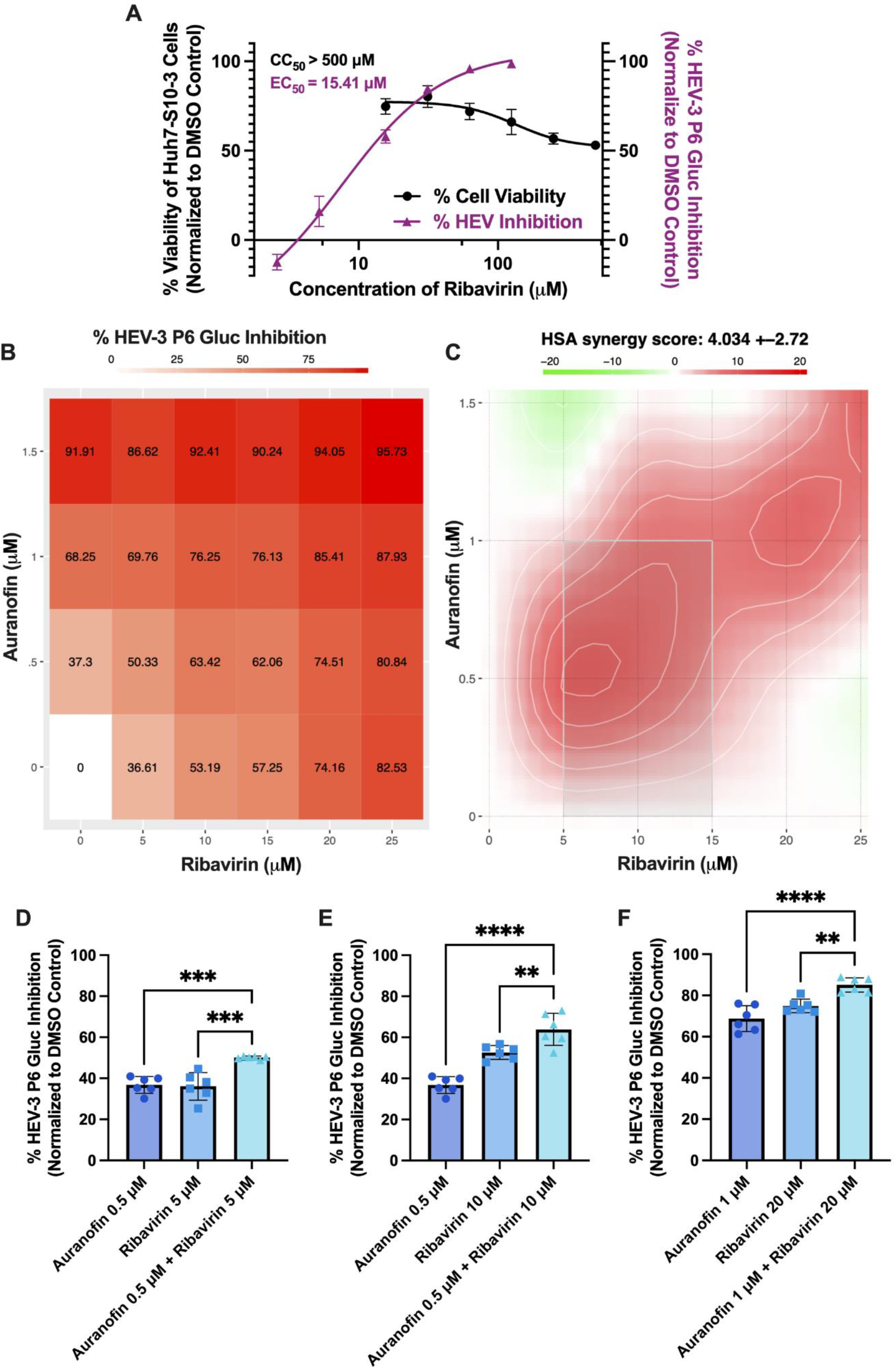
Auranofin and ribavirin combined treatment *in vitro* displays synergistic antiviral activity. (*A*) The effect of ribavirin on cytotoxicity of Huh7-S10-3 cells and antiviral activity against the HEV-3 P6 Gluc replicon. Data collected 72 hours post compound application is background subtracted and normalized to the vehicle control. X-axis values are log transformed. The lines represent non-linear regression curves, and data points represent means ± SD; n=3 performed in two independent experiments. (*B & C*) Huh7-S10-3 cells were transfected with HEV-3 P6 Gluc RNA, compounds were applied, and Gluc was quantified from supernatants 72 hours post complication. Resulting data was generated via the SynergyFinder web application (version 3.0). (*B*) Dose-response HEV-3 P6 Gluc inhibition matrix of the combined concentrations of auranofin and ribavirin tested at a range between 0 - 1.5 μM and 0 - 25 μM, respectively. The average inhibition of HEV replication is presented for all tested compound combinations and for auranofin and ribavirin treatment alone. (*C*) Synergy map generated via the HSA model. The overall HSA synergy score is depicted as 4.034 ± 2.72. (*D-F*) Graphs depicting the concentration combinations that displayed synergy (synergy scores over 10): (*D*) auranofin 0.5 μM + ribavirin 5 μM, *(E*) auranofin 0.5 μM + ribavirin 10 μM, and (*F*) auranofin 1 μM + ribavirin 20 μM. Data in B-F is background subtracted and normalized to the appropriate vehicle control. A one-way ANOVA test was performed for D-F. The data points represent means ± SD; n=6, performed in two independent experiments.

To determine the potential synergistic antiviral activity, auranofin and ribavirin were tested against HEV-3 P6 Gluc at concentrations in combination ranging from 0 - 1.5 μM and 0 - 25 μM, respectively. Antiviral data were uploaded to the SynergyFinder web application (version 3.0) to identify potential synergy, defined as a synergy score greater than 10. The highest single agent (HSA) model was used, which determines whether the combined effect of the two drugs exceeds the sum of their individual effects. This provides a stronger weight towards lower concentrations, which are less likely to display toxicity (59). We observed a dose-response to combined auranofin and ribavirin treatments against HEV-3 P6 Gluc for each of the tested concentration combinations (Fig. 4B), with average inhibition scores presented. Based on the HSA model, 3 of the concentration combinations tested in the study display a synergy score over 10, which can be interpreted as 10% of response beyond the individual compound effects (59) (Supplementary File 1), resulting in an overall HSA synergy score of 4.034 ± 2.72 (Fig. 4C). The 3 combinations that display synergy are: 0.5 μM auranofin + 5 μM ribavirin with a synergy score of 13.03 (Fig. 4D), 0.5 μM auranofin + 10 μM ribavirin with a synergy score of 10.235 (Fig. 4E), and 1 μM auranofin + 20 μM ribavirin with a synergy score of 11.255 (Fig. 4F). Overall, this data demonstrates that auranofin and ribavirin display synergistic antiviral activity at select concentrations *in vitro*.

## Discussion

HEV infection is associated with serious clinical conditions, such as chronic infection, fulminant hepatic failure, and neurological sequelae that require effective antiviral intervention. Due to the emergence of HEV strains resistant to ribavirin (37, 38), the current off-label standard of care, it is essential to identify HEV-specific and more effective antivirals that act via different mechanisms. Because auranofin is FDA-approved and reported to be an effective antiviral against several other viruses (46, 47, 71), we investigated the potential of auranofin as an antiviral against HEV. We demonstrated that auranofin displays dose-dependent antiviral activity against two different genotypes of HEV (HEV-1 and HEV-3) and a HEV-3 mutant associated with ribavirin treatment failure. We further showed that select concentrations of auranofin and ribavirin in combined treatments display synergistic antiviral activity, supporting a combined therapeutic strategy to enhance antiviral efficacy and prevent resistance to either drug. Further work to parse out the mechanism of action revealed a causal role for ROS in driving the antiviral activity of auranofin. Consistent with this observation, we also showed that ROS induced by an independent mechanism displayed antiviral activity, highlighting the promotion of ROS as a novel antiviral strategy against HEV.

Auranofin’s anticancer activity has been linked to the accumulation of ROS via the inhibition of the redox enzymes, GPx (72) and TrxR (73), that play important roles in the glutathione and thioredoxin redox systems, respectively. These disulfide reductase systems are antioxidant in nature and function to maintain ROS homeostasis within cells (74). Numerous studies present the connection between the inhibition of these enzymes, the upregulation of ROS, and auranofin’s anticancer effects (62, 75, 76). This connection is supported by the finding that NAC, a ROS inhibitor, and auranofin co-treatment reverse the toxicity of auranofin (63, 75, 76). Here, we found that NAC also reversed auranofin’s antiviral activity, a novel finding in the investigation of auranofin’s mechanism of action against HEV.

While several studies have previously reported auranofin’s antiviral activity against other viruses, different mechanisms of antiviral action have been suggested. Auranofin was shown to be an effective antiviral against chikungunya virus *in vivo*; however, the mechanism of action was not investigated (71). Auranofin was also reported to impact the viral reservoir of human immunodeficiency virus type 1 (HIV-1); auranofin induced a pro-apoptotic effect via a burst of reactive oxygen species (ROS) in CD4+ T cells, which are typically unaffected by antiretroviral therapy (ART) (46). During a randomized clinical trial, auranofin in conjunction with ART was able to decrease total integrated HIV-1 DNA compared to ART alone, suggesting its efficacy against HIV-1 infection in humans (45). Auranofin was also found to inhibit SARS-CoV-2 replication and reduce cytokine production, which supports a dual action effect on the virus and associated lung injury and disease severity induced by cytokine storms (47). Auranofin and similar analogs have also been linked to anti-protease activity against SARS-CoV-2 (48). While these studies suggest differential mechanisms for auranofin’s apparent broad-spectrum antiviral activity, the modification of redox pathways is a common characteristic discussed, suggesting that modulation of the redox pathways contributes to auranofin’s antiviral effects. Our work with auranofin and HEV provides further support for this connection and may be the common underlying mechanism driving auranofin’s antiviral activity.

While repurposing auranofin as a broad-spectrum antiviral is well-supported, more work is needed to fully elucidate its antiviral mechanism(s) and its connection to redox modulation and ROS promotion. Many viruses modulate ROS and ROS signaling pathways; thus, modulating ROS has been suggested as a potential antiviral target (77, 78). While too much ROS can be detrimental to cells, inducing apoptosis, an increase in ROS at non-toxic levels has been shown to activate signaling pathways for various cellular processes. Of interest are pathways related to antioxidants, inflammation, antiviral immunity, and DNA damage (79). Through further investigations into the pathways activated by ROS promotion, it is possible to identify new antiviral targets that may bypass ROS-induced effects and potential cytotoxicity, leading to targeted and safer antiviral therapeutics.

More work is also needed to explore auranofin’s specific antiviral effects against HEV. While we validated the *in vitro* antiviral effects of auranofin against two important genotypes of HEV (HEV-1 and HEV-3), which are responsible for significant proportions of human disease, it is also important in the future to further validate auranofin’s antiviral efficacy against HEV using an *in vivo* model, which is beyond the scope of this current study. This represents a challenge as immunocompetent small animal models of HEV infection are available, but working with them requires overcoming obstacles related to genotype specificity, lack of recapitulated disease, and high costs associated with animal care and housing (80). However, further investigation into the *in vivo* efficacy of auranofin as a stand-alone therapeutic against HEV, a combined treatment with ribavirin, and as a treatment against ribavirin resistant HEV infections is warranted. Further work with auranofin could also explore antiviral efficacy against chronic HEV infections, as ribavirin resistance arises in individuals with chronic HEV infections and liver failure (81). However, while more work is needed to further validate auranofin’s antiviral efficacy *in vivo*, our current *in vitro* data does support the repurposing of auranofin as an antiviral against HEV, therefore laying the foundation for further research and development of a HEV-specific antiviral which is currently lacking.

In conclusion, we have identified dose-dependent antiviral effects *in vitro* of auranofin against two different genotypes of HEV and a HEV mutant associated with ribavirin treatment failure. These antiviral effects are linked to ROS, as ROS inhibitors reverse the antiviral effects and ROS promotion caused by auranofin treatment. To further validate the connection between ROS and antiviral activity, we tested the ROS promotor DAAO, which also displayed strong antiviral activity, which was reversed in the presence of NAC along with ROS promotion. This further highlights ROS promotion as a novel antiviral target against HEV, as DAAO acts through an alternative mechanism via the breakdown of D-amino acids. A combined treatment strategy of auranofin and ribavirin *in vitro* suggests that a synergistic antiviral activity can be achieved at select concentrations, which provides combinational drug therapeutic potential to increase antiviral activity and minimize ribavirin resistance. Altogether, this data supports our conclusion that auranofin inhibits HEV replication, driven by redox modulations and the promotion of ROS. This finding is novel and significant because HEV infections can become resistant to ribavirin, the current standard of care via off-label use, and there are no approved HEV-specific therapeutics.Therefore, repurposing auranofin could provide more treatment options for those experiencing chronic and severe HEV infections. Overall, our *in vitro* data is supportive of further testing of auranofin *in vivo* against HEV infections and further testing to explore the connection between ROS upregulation and antiviral activity.

## Acknowledgements

We thank Drs Suzanne U. Emerson and Robert H. Purcell (NIAID, NIH, Bethesda, MD) for providing the Huh7-S10-3 cell line, infectious clones of HEV genotype 3 Kernow-C1 strain (P6) and HEV genotype 1 Sar55 strain, and HEV-3 P6 Gluc indicator replicon. We also thank Dr. Alexander Ploss (Princeton University, Princeton, NJ) for providing the HEV-1 Sar55 Gluc indicator replicon. We are grateful to Melissa Makris for analysis of flow cytometry data. This work was supported by a grant from the National Science Foundation through the Predictive Intelligence for Pandemic Prevention (PIPP) program awarded to JWL and XJM. Additional support was provided in part by a seed grant from the Virginia Tech Center for Drug Discovery (VTCDD, to JWL) and grants (R01AI050611, R37AI179614 to XJM) from the National Institutes of Health.

## References

1. WORLD HEALTH ORGANIZATION. 2015. Hepatitis E vaccine: WHOposition paper, May 2015. Wkly Epidemiol Rec.

2. Rein DB, Stevens GA, Theaker J, Wittenborn JS, Wiersma ST. 2012. The global burden of hepatitis E virus genotypes 1 and 2 in 2005. Hepatology 55:988–997.

3. Webb GW, Dalton HR. 2019. Hepatitis E: an underestimated emerging threat. Ther Adv Infect Dis 6:2049936119837162.

4. Dalton HR, Kamar N, Izopet J. 2014. Hepatitis E in developed countries: current status and future perspectives. Future Microbiol 9:1361–1372.

5. Meng X-J. 2013. Zoonotic and foodborne transmission of hepatitis E virus. Semin Liver Dis 33:41–49.

6. Hakim MS, Wang W, Bramer WM, Geng J, Huang F, de Man RA, Peppelenbosch MP, Pan Q. 2017. The global burden of hepatitis E outbreaks: a systematic review. Liver Int 37:19–31.

7. European Association for the Study of the Liver. 2018. EASL Clinical Practice Guidelines on hepatitis E virus infection. J Hepatol 68:1256–1271.

8. Purdy MA, Drexler JF, Meng X-J, Norder H, Okamoto H, Van der Poel WHM, Reuter G, de Souza WM, Ulrich RG, Smith DB. 2022. ICTV virus Taxonomy profile: Hepeviridae 2022. J Gen Virol 103.

9. Wang B, Meng X-J. 2021. Hepatitis E virus: host tropism and zoonotic infection. Curr Opin Microbiol 59:8–15.

10. Balayan MS, Andjaparidze AG, Savinskaya SS, Ketiladze ES, Braginsky DM, Savinov AP, Poleschuk VF. 1983. Evidence for a virus in non-A, non-B hepatitis transmitted via the fecal-oral route. Intervirology 20:23–31.

11. Nagashima S, Takahashi M, Kobayashi T, Tanggis Nishizawa T, Nishiyama T, Primadharsini PP, Okamoto H. 2017. Characterization of the quasi-enveloped hepatitis E virus particles released by the cellular exosomal pathway. J Virol 91.

12. Takahashi M, Yamada K, Hoshino Y, Takahashi H, Ichiyama K, Tanaka T, Okamoto H. 2008. Monoclonal antibodies raised against the ORF3 protein of hepatitis E virus (HEV) can capture HEV particles in culture supernatant and serum but not those in feces. Arch Virol 153:1703–1713.

13. Yin X, Ambardekar C, Lu Y, Feng Z. 2016. Distinct entry mechanisms for nonenveloped and quasi-enveloped hepatitis E viruses. J Virol 90:4232–4242.

14. Tam AW, Smith MM, Guerra ME, Huang CC, Bradley DW, Fry KE, Reyes GR. 1991. Hepatitis E virus (HEV): molecular cloning and sequencing of the full-length viral genome. Virology 185:120–131.

15. Kenney SP, Meng X-J. 2019. Hepatitis E Virus Genome Structure and Replication Strategy. Cold Spring Harb Perspect Med 9.

16. Yin X, Ying D, Lhomme S, Tang Z, Walker CM, Xia N, Zheng Z, Feng Z. 2018. Origin, antigenicity, and function of a secreted form of ORF2 in hepatitis E virus infection. Proc Natl Acad Sci U S A 115:4773–4778.

17. Ding Q, Heller B, Capuccino JMV, Song B, Nimgaonkar I, Hrebikova G, Contreras JE, Ploss A. 2017. Hepatitis E virus ORF3 is a functional ion channel required for release of infectious particles. Proc Natl Acad Sci U S A 114:1147–1152.

18. Kumar Acharya S, Kumar Sharma P, Singh R, Kumar Mohanty S, Madan K, Kumar Jha J, Kumar Panda S. 2007. Hepatitis E virus (HEV) infection in patients with cirrhosis is associated with rapid decompensation and death. J Hepatol 46:387–394.

19. Kumar A, Beniwal M, Kar P, Sharma JB, Murthy NS. 2004. Hepatitis E in pregnancy. Int J Gynaecol Obstet 85:240–244.

20. Buescher G, Ozga A-K, Lorenz E, Pischke S, May J, Addo MM, Horvatits T. 2021. Hepatitis E seroprevalence and viremia rate in immunocompromised patients: a systematic review and meta-analysis. Liver Int 41:449–455.

21. Dalton HR, Bendall RP, Keane FE, Tedder RS, Ijaz S. 2009. Persistent carriage of hepatitis E virus in patients with HIV infection. N Engl J Med 361:1025–1027.

22. Kamar N, Mansuy J-M, Cointault O, Selves J, Abravanel F, Danjoux M, Otal P, Esposito L, Durand D, Izopet J, Rostaing L. 2008. Hepatitis E virus-related cirrhosis in kidney- and kidney-pancreas-transplant recipients. Am J Transplant 8:1744– 1748.

23. Geng Y, Zhang H, Huang W. J Harrison T, Geng K, Li Z, Wang Y. 2014. Persistent hepatitis e virus genotype 4 infection in a child with acute lymphoblastic leukemia. Hepat Mon 14:e15618.

24. Jilani N, Das BC, Husain SA, Baweja UK, Chattopadhya D, Gupta RK, Sardana S, Kar P. 2007. Hepatitis E virus infection and fulminant hepatic failure during pregnancy: Hepatitis E virus and fulminant hepatic failure. J Gastroenterol Hepatol 22:676–682.

25. Qian Z, Li T, Zhang Y, Chen S, Zhang H, Mickael HK, Xiu D, Xia Y, Cong C, Xu L, Wei D, Yu W, Yu X, Huang F. 2023. Prevalence of hepatitis E virus and its association with adverse pregnancy outcomes in pregnant women in China. J Clin Virol 158:105353.

26. Tsarev SA, Tsareva TS, Emerson SU, Rippy MK, Zack P, Shapiro M, Purcell RH. 1995. Experimental hepatitis E in pregnant rhesus monkeys: failure to transmit hepatitis E virus (HEV) to offspring and evidence of naturally acquired antibodies to HEV. J Infect Dis 172:31–37.

27. Guo H, Zhou EM, Sun ZF, Meng X-J. 2007. Egg whites from eggs of chickens infected experimentally with avian hepatitis E virus contain infectious virus, but evidence of complete vertical transmission is lacking. J Gen Virol 88:1532–1537.

28. Ripellino P, Pasi E, Melli G, Staedler C, Fraga M, Moradpour D, Sahli R, Aubert V, Martinetti G, Bihl F, Bernasconi E, Terziroli Beretta-Piccoli B, Cerny A, Dalton HR, Zehnder C, Mathis B, Zecca C, Disanto G, Kaelin-Lang A, Gobbi C. 2020. Neurologic complications of acute hepatitis E virus infection. Neurol Neuroimmunol Neuroinflamm 7:e643.

29. Kamar N, Bendall RP, Peron JM, Cintas P, Prudhomme L, Mansuy JM, Rostaing L, Keane F, Ijaz S, Izopet J, Dalton HR. 2011. Hepatitis E virus and neurologic disorders. Emerg Infect Dis 17:173–179.

30. van den Berg B, van der Eijk AA, Pas SD, Hunter JG, Madden RG, Tio-Gillen AP, Dalton HR, Jacobs BC. 2014. Guillain-Barré syndrome associated with preceding hepatitis E virus infection. Neurology 82:491–497.

31. Food and Agriculture Organization of the United Nations, World Health Organization. 2023. Joint FAO/WHO Expert Meeting on microbiological risk assessment of viruses in food Part 1: food attribution, analytical methods, and indicators.

32. Ji Y, Li P, Jia Y, Wang X, Zheng Q, Peppelenbosch MP, Ma Z, Pan Q. 2021. Estimating the burden and modeling mitigation strategies of pork-related hepatitis E virus foodborne transmission in representative European countries. One Health 13:100350.

33. García N, Hernández M, Gutierrez-Boada M, Valero A, Navarro A, Muñoz-Chimeno M, Fernández-Manzano A, Escobar FM, Martínez I, Bárcena C, González S, Avellón A, Eiros JM, Fongaro G, Domínguez L, Goyache J, Rodríguez-Lázaro D. 2019. Occurrence of hepatitis E virus in pigs and pork cuts and organs at the time of slaughter, Spain, 2017. Front Microbiol 10:2990.

34. Renou C, Roque-Afonso A-M, Pavio N. 2014. Foodborne transmission of hepatitis E virus from raw pork liver sausage, France. Emerg Infect Dis 20:1945–1947.

35. Zhu F-C, Zhang J, Zhang X-F, Zhou C, Wang Z-Z, Huang S-J, Wang H, Yang C-L, Jiang H-M, Cai J-P, Wang Y-J, Ai X, Hu Y-M, Tang Q, Yao X, Yan Q, Xian Y-L, Wu T, Li Y-M, Miao J, Ng M-H, Shih JW-K, Xia N-S. 2010. Efficacy and safety of a recombinant hepatitis E vaccine in healthy adults: a large-scale, randomised, double-blind placebo-controlled, phase 3 trial. Lancet 376:895–902.

36. Hartley C, Wasuwanich P, Van T, Karnsakul W. 2024. Hepatitis E vaccines updates. Vaccines (Basel) 12:722.

37. Debing Y, Ramière C, Dallmeier K, Piorkowski G, Trabaud M-A, Lebossé F, Scholtès C, Roche M, Legras-Lachuer C, de Lamballerie X, André P, Neyts J. 2016. Hepatitis E virus mutations associated with ribavirin treatment failure result in altered viral fitness and ribavirin sensitivity. J Hepatol 65:499–508.

38. Wang B, Mahsoub HM, Li W, Heffron CL, Tian D, Hassebroek AM, LeRoith T, Meng X-J. 2023. Ribavirin treatment failure-associated mutation, Y1320H, in the RNA-dependent RNA polymerase of genotype 3 hepatitis E virus (HEV) enhances virus replication in a rabbit HEV infection model. MBio 14:e0337222.

39. Puoti M, Babudieri S, Rezza G, Viale P, Antonini MG, Maida I, Rossi S, Zanini B, Putzolu V, Fenu L, Baiguera C, Sassu S, Carosi G, Mura MS. 2004. Use of pegylated interferons is associated with an increased incidence of infections during combination treatment of chronic hepatitis C: a side effect of pegylation? Antivir Ther 9:627–630.

40. Kowdley K. 2005. Hematologic side effects of interferon and ribavirin therapy. J Clin Gastroenterol 39:S3–8.

41. Sulkowski MS, Cooper C, Hunyady B, Jia J, Ogurtsov P, Peck-Radosavljevic M, Shiffman ML, Yurdaydin C, Dalgard O. 2011. Management of adverse effects of Peg-IFN and ribavirin therapy for hepatitis C. Nat Rev Gastroenterol Hepatol 8:212–223.

42. Martinez MA. 2022. Efficacy of repurposed antiviral drugs: Lessons from COVID-19. Drug Discov Today 27:1954–1960.

43. Trivedi J, Mohan M, Byrareddy SN. 2020. Drug repurposing approaches to combating viral infections. J Clin Med 9:3777.

44. Shen S, Shen J, Luo Z, Wang F, Min J. 2023. Molecular mechanisms and clinical implications of the gold drug auranofin. Coord Chem Rev 493:215323.

45. Diaz RS, Shytaj IL, Giron LB, Obermaier B, Della Libera E Jr, Galinskas J, Dias D, Hunter J, Janini M, Gosuen G, Ferreira PA, Sucupira MC, Maricato J, Fackler O, Lusic M, Savarino A, SPARC Working Group. 2019. Potential impact of the antirheumatic agent auranofin on proviral HIV-1 DNA in individuals under intensified antiretroviral therapy: Results from a randomised clinical trial. Int J Antimicrob Agents 54:592–600.

46. Chirullo B, Sgarbanti R, Limongi D, Shytaj IL, Alvarez D, Das B, Boe A, DaFonseca S, Chomont N, Liotta L, Petricoin E Iii, Norelli S, Pelosi E, Garaci E, Savarino A, Palamara AT. 2013. A candidate anti-HIV reservoir compound, auranofin, exerts a selective “anti-memory” effect by exploiting the baseline oxidative status of lymphocytes. Cell Death Dis 4:e944.

47. Rothan HA, Stone S, Natekar J, Kumari P, Arora K, Kumar M. 2020. The FDA-approved gold drug auranofin inhibits novel coronavirus (SARS-COV-2) replication and attenuates inflammation in human cells. Virology 547:7–11.

48. Massai L, Grifagni D, De Santis A, Geri A, Cantini F, Calderone V, Banci L, Messori L. 2022. Gold-Based Metal Drugs as Inhibitors of Coronavirus Proteins: The Inhibition of SARS-CoV-2 Main Protease by Auranofin and Its Analogs. Biomolecules 12.

49. Emerson SU, Nguyen H, Torian U, Purcell RH. 2006. ORF3 protein of hepatitis E virus is not required for replication, virion assembly, or infection of hepatoma cells in vitro. J Virol 80:10457–10464.

50. Nguyen HT, Shukla P, Torian U, Faulk K, Emerson SU. 2014. Hepatitis E virus genotype 1 infection of swine kidney cells in vitro is inhibited at multiple levels. J Virol 88:868–877.

51. Ding Q, Nimgaonkar I, Archer NF, Bram Y, Heller B, Schwartz RE, Ploss A. 2018. Identification of the intragenomic promoter controlling hepatitis E virus subgenomic RNA transcription. MBio 9.

52. Shukla P, Nguyen HT, Torian U, Engle RE, Faulk K, Dalton HR, Bendall RP, Keane FE, Purcell RH, Emerson SU. 2011. Cross-species infections of cultured cells by hepatitis E virus and discovery of an infectious virus-host recombinant. Proc Natl Acad Sci U S A 108:2438–2443.

53. Shukla P, Nguyen HT, Faulk K, Mather K, Torian U, Engle RE, Emerson SU. 2012. Adaptation of a genotype 3 hepatitis E virus to efficient growth in cell culture depends on an inserted human gene segment acquired by recombination. J Virol 86:5697–5707.

54. Debing Y, Gisa A, Dallmeier K, Pischke S, Bremer B, Manns M, Wedemeyer H, Suneetha PV, Neyts J. 2014. A mutation in the hepatitis E virus RNA polymerase promotes its replication and associates with ribavirin treatment failure in organ transplant recipients. Gastroenterology 147:1008–11.e7; quiz e15–6.

55. Wang B, Tian D, Sooryanarain H, Mahsoub HM, Heffron CL, Hassebroek AM, Meng X-J. 2022. Two mutations in the ORF1 of genotype 1 hepatitis E virus enhance virus replication and may associate with fulminant hepatic failure. Proc Natl Acad Sci U S A 119:e2207503119.

56. Tian D, Li W, Heffron LC, Mahsoub HM, Wang B, LeRoith T, and Meng XJ. 2025. Antiviral resistance and barrier integrity at the maternal-fetal interface restricts hepatitis E virus from crossing the placental barrier. Proc Natl Acad Sci U S A 10.1073/pnas.2501128122.

57. Marano JM, Chuong C, Weger-Lucarelli J. 2020. Rolling circle amplification: A high fidelity and efficient alternative to plasmid preparation for the rescue of infectious clones. Virology 551:58–63.

58. Vargas MH. 2000. ED50plus (v1.0). https://archive.org/details/ed50v10_zip.

59. Ianevski A, Giri AK, Aittokallio T. 2022. SynergyFinder 3.0: an interactive analysis and consensus interpretation of multi-drug synergies across multiple samples. Nucleic Acids Res 50:W739–W743.

60. Radenkovic F, Holland O, Vanderlelie JJ, Perkins AV. 2017. Selective inhibition of endogenous antioxidants with Auranofin causes mitochondrial oxidative stress which can be countered by selenium supplementation. Biochem Pharmacol 146:42–52.

61. Wang H, Bouzakoura S, de Mey S, Jiang H, Law K, Dufait I, Corbet C, Verovski V, Gevaert T, Feron O, Van den Berge D, Storme G, De Ridder M. 2017. Auranofin radiosensitizes tumor cells through targeting thioredoxin reductase and resulting overproduction of reactive oxygen species. Oncotarget 8:35728–35742.

62. Park S-J, Kim I-S. 2005. The role of p38 MAPK activation in auranofin-induced apoptosis of human promyelocytic leukaemia HL-60 cells. Br J Pharmacol 146:506–513.

63. Cui XY, Park SH, Park WH. 2022. Anti-Cancer Effects of Auranofin in Human Lung Cancer Cells by Increasing Intracellular ROS Levels and Depleting GSH Levels. Molecules 27.

64. Aldini G, Altomare A, Baron G, Vistoli G, Carini M, Borsani L, Sergio F. 2018. N-Acetylcysteine as an antioxidant and disulphide breaking agent: the reasons why. Free Radic Res 52:1–12.

65. Zafarullah M, Li WQ, Sylvester J, Ahmad M. 2003. Molecular mechanisms of N-acetylcysteine actions. Cell Mol Life Sci 60:6–20.

66. Rodrigues RB, de Oliveira MM, Garcia FP, Ueda-Nakamura T, de Oliveira Silva S, Nakamura CV. 2024. Dithiothreitol reduces oxidative stress and necrosis caused by ultraviolet A radiation in L929 fibroblasts. Photochem Photobiol Sci 23:271–284.

67. Ma W-X, Li C-Y, Tao R, Wang X-P, Yan L-J. 2020. Reductive stress-induced mitochondrial dysfunction and cardiomyopathy. Oxid Med Cell Longev 2020:5136957.

68. Spyropoulos F, Michel T. 2024. D-Amino acid oxidase-derived chemogenetic oxidative stress: Unraveling the multi-omic responses to in vivo redox stress. Curr Opin Chem Biol 79:102438.

69. Debing Y, Emerson SU, Wang Y, Pan Q, Balzarini J, Dallmeier K, Neyts J. 2014. Ribavirin inhibits in vitro hepatitis E virus replication through depletion of cellular GTP pools and is moderately synergistic with alpha interferon. Antimicrob Agents Chemother 58:267–273.

70. Marimani M, Ahmad A, Duse A. 2020. Combination therapy as an effective tool for treatment of drug-resistant viral infections, p. 157–182. In Combination Therapy Against Multidrug Resistance. Elsevier.

71. Langsjoen RM, Auguste AJ, Rossi SL, Roundy CM, Penate HN, Kastis M, Schnizlein MK, L. KC, Haller SL, Chen R, Watowich SJ, Weaver SC. 2017. Host oxidative folding pathways offer novel anti-chikungunya virus drug targets with broad spectrum potential. Antiviral Res 143:246–251.

72. Kudin AP, Augustynek B, Lehmann AK, Kovács R, Kunz WS. 2012. The contribution of thioredoxin-2 reductase and glutathione peroxidase to H(2)O(2) detoxification of rat brain mitochondria. Biochim Biophys Acta 1817:1901–1906.

73. Marzano C, Gandin V, Folda A, Scutari G, Bindoli A, Rigobello MP. 2007. Inhibition of thioredoxin reductase by auranofin induces apoptosis in cisplatin-resistant human ovarian cancer cells. Free Radic Biol Med 42:872–881.

74. Ren X, Zou L, Zhang X, Branco V, Wang J, Carvalho C, Holmgren A, Lu J. 2017. Redox Signaling Mediated by Thioredoxin and Glutathione Systems in the Central Nervous System. Antioxid Redox Signal 27:989–1010.

75. You BR, Park WH. 2016. Auranofin induces mesothelioma cell death through oxidative stress and GSH depletion. Oncol Rep 35:546–551.

76. Abdalbari FH, Martinez-Jaramillo E, Forgie BN, Tran E, Zorychta E, Goyeneche AA, Sabri S, Telleria CM. 2023. Auranofin Induces Lethality Driven by Reactive Oxygen Species in High-Grade Serous Ovarian Cancer Cells. Cancers 15.

77. Sander WJ, Fourie C, Sabiu S, O’Neill FH, Pohl CH, O’Neill HG. 2022. Reactive oxygen species as potential antiviral targets. Rev Med Virol 32:e2240.

78. Nencioni L, Sgarbanti R, Amatore D, Checconi P, Celestino I, Limongi D, Anticoli S, Palamara AT, Garaci E. 2011. Intracellular redox signaling as therapeutic target for novel antiviral strategy. Curr Pharm Des 17:3898–3904.

79. Li Z, Xu X, Leng X, He M, Wang J, Cheng S, Wu H. 2017. Roles of reactive oxygen species in cell signaling pathways and immune responses to viral infections. Arch Virol 162:603–610.

80. Xiang Z, He X-L, Zhu C-W, Yang J-J, Huang L, Jiang C, Wu J, Chinese Consortium for the Study of Hepatitis E (CCSHE). 2024. Animal models of hepatitis E infection: Advances and challenges. Hepatobiliary Pancreat Dis Int 23:171–180.

81. Kamar N, Izopet J, Pavio N, Aggarwal R, Labrique A, Wedemeyer H, Dalton HR. 2017. Hepatitis E virus infection. Nat Rev Dis Primers 3:17086.

